# The *Escherichia coli serS* gene promoter region overlaps with the *rarA* gene

**DOI:** 10.1101/2021.11.08.467797

**Authors:** Kanika Jain, Tyler H. Stanage, Elizabeth A. Wood, Michael M. Cox

## Abstract

Deletion of the entire gene encoding the RarA protein of *Escherichia coli* results in a growth defect and additional deficiencies that were initially ascribed to a lack of RarA function. Further work revealed that most of the effects reflected the presence of sequences in the *rarA* gene that affect expression of the downstream gene, *serS.* The *serS* gene encodes the seryl aminoacyl-tRNA synthetase. Decreases in the expression of *serS* can trigger the stringent response. The sequences that affect *serS* expression are located in the last 15 nucleotides of the *rarA* gene.

## Introduction

When a replication fork encounters roadblocks, such as DNA lesions, template strand breaks, or DNA-bound proteins, it can stall. Outcomes may include fork collapse and replisome dissociation (1–11). These events can have catastrophic consequences for genomic integrity and cell viability, if left unrepaired. In bacteria, estimates vary, but replication forks may stall as often as once per cell generation during normal growth conditions (2, 12–20). Most of the adverse replication fork encounters are resolved using a variety of pathways that do not introduce mutations (2, 3, 7, 9–13, 21–26). Sometimes, a fork skips over the lesion and re-initiates downstream, leaving the lesion behind in what is called as a post-replication gap (4, 8, 13, 27–31). There appear to be three major paths for filling post-replication gaps in bacteria: (a) RecA-mediated homologous recombination (32–35), (b) translesion DNA synthesis (1, 36, 37), and (c) a RecA-independent template switching process (38–41).The *Escherichia coli* RarA protein is required for most of this RecA-independent recombination (41). More prominently, the RarA protein is involved in the resolution of recombination intermediates as part of an expanded RecFOR pathway for the amelioration of post-replication gaps (*manuscript under review*).

The *Escherichia coli* RarA protein is an ATPase in the AAA+ superfamily (42, 43). The *rarA* gene encodes a 447-amino-acid polypeptide with a predicted monomeric mass of 49594 kDa. The protein is part of a highly conserved family. It is absent in archaea but highly conserved from bacteria through eukaryotes, sharing about 40% identity and 56-58% similarity with its *Saccharomyces cerevisiae* (Mgs1) and Homo sapiens (WRNIP1) homologs (42, 43). In *E. coli,* RarA shares 25% amino acid identity with two other proteins: RuvB, a Holliday Junction helicase, and DnaX, a subunit of DNA polymerase III replisome. DnaX encodes for Tau (τ) and Gamma (γ) components of DNA polymerase III clamp loader complex, placing RarA in the clamp loader AAA+ clade (42, 43).

In *E. coli* chromosome, the *rarA* gene is located at 20.21 centisomes (location 937,994>939,337). The *rarA* gene is immediately upstream of an essential gene, *serS*, a serine-tRNA ligase, located at 20.24 centisomes. SerS is among the 20 aminoacyl-tRNA synthetases (aaRSs) or tRNA-ligases present in the cell. aaRSs are the charging portals of tRNAs. They generate a covalent linkage between an amino acid and its cognate tRNA to form an aminoacyl-tRNA complex. The ribosome acts on this charged tRNA complex and transfers its attached amino acid onto the growing peptide chain - thereby fostering the translation process in the cell. SerS aminoacylates tRNA^Ser^ and tRNA^Sec^ with serine (44, 45). *serS* is mainly regulated by its promoter (serSp1) with a transcription start located at 939,365^th^ position after the end of *rarA* gene (46) (**Figure 1**).

**Figure 1:**
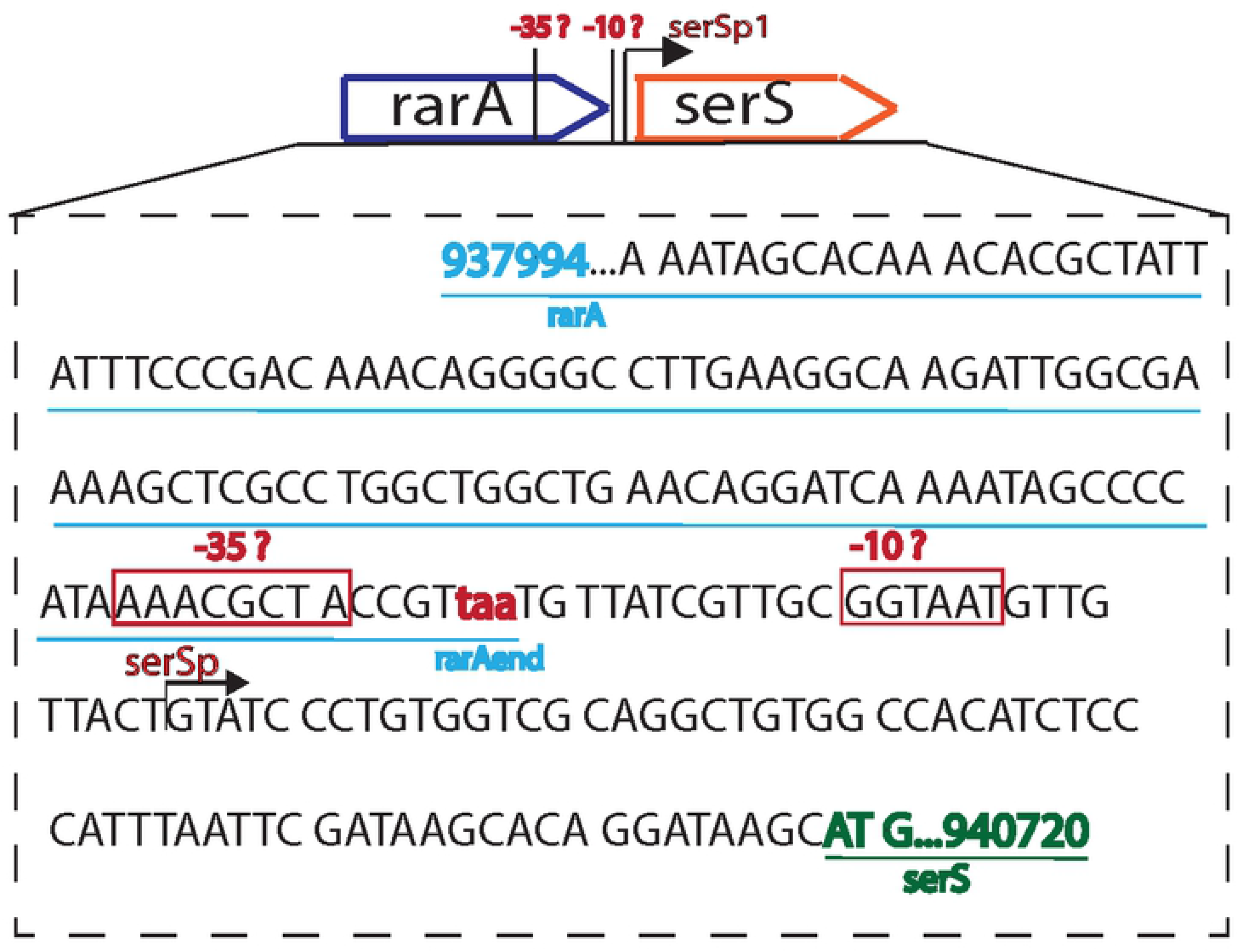
Identification of possible promoter/regulatory sequence of *serS* within the *rarA* sequence. Representation of predicted promoter sequences and their location in the last 40 amino acids region of *rarA.*

aaRSs manage the growth and the stringent response in the cell by directly controlling the two interdependent cellular processes: (1) the flux of protein synthesis, and (2) the levels of uncharged tRNAs. The first process - the flux of protein synthesis - is directly dependent on the amount of tRNA aminoacylated by aaRSs. Modifications in aaRSs production impedes cell growth (47, 48). Globally, high levels of uncharged tRNA slow translation kinetics and thereby slow cell growth - both in bacteria and eukaryotes (49–51). In bacteria, these high levels of uncharged tRNA are detected by the (p)ppGpp synthetase -RelA- which in response induces a stringent response and affects the cell growth (52–54).

SerS is notable as it is inhibited by serine hydroxamate, a small molecule often used by investigators to induce the stringent response (55). We have found that the complete deletion of the *rarA* gene slows cell growth, impedes SOS induction, and rescues DNA damage sensitivity of several repair-deficient cells, effects we initially attributed to a lack of RarA function. This initial conclusion was in error. All of these phenotypes disappear when a slightly more modest *rarA* deletion is used that deletes more than 90% of the coding sequence, all but the last 41 codons. This suggests that regulatory sequences that affect *serS* expression may be embedded in the *rarA* coding sequence. Keeping a small portion of the *rarA* gene, that which encodes C-terminal of RarA, is vital for optimal growth of the *E. coli* cell. A –35 segment of the *serS* promoter or some equivalent regulatory sequence appears to be located in the last 15 nucleotides of the *rarA* gene.

## Materials and Methods

### Strain construction

All strains are *E. coli* MG1655 derivatives and are listed in **Table 1.** Some of the *rarA* strains (*rarA*N406, *rarA*N430, *rarA*N437 and *rarA*N442) were made using *galK+* recombineering method. *rarA*Δ*N447* (EAW98) and all other strains were constructed using Lambda red recombination as described by Datsenko and Wanner (59). Kanamycin resistance of these strains was removed using FLP recombinase when required (62). All chromosomal mutations were confirmed using Sanger sequencing. Standard transformation protocols were used to generate strains harboring the indicated plasmids as listed in **Table 1**.

**Table 1:**
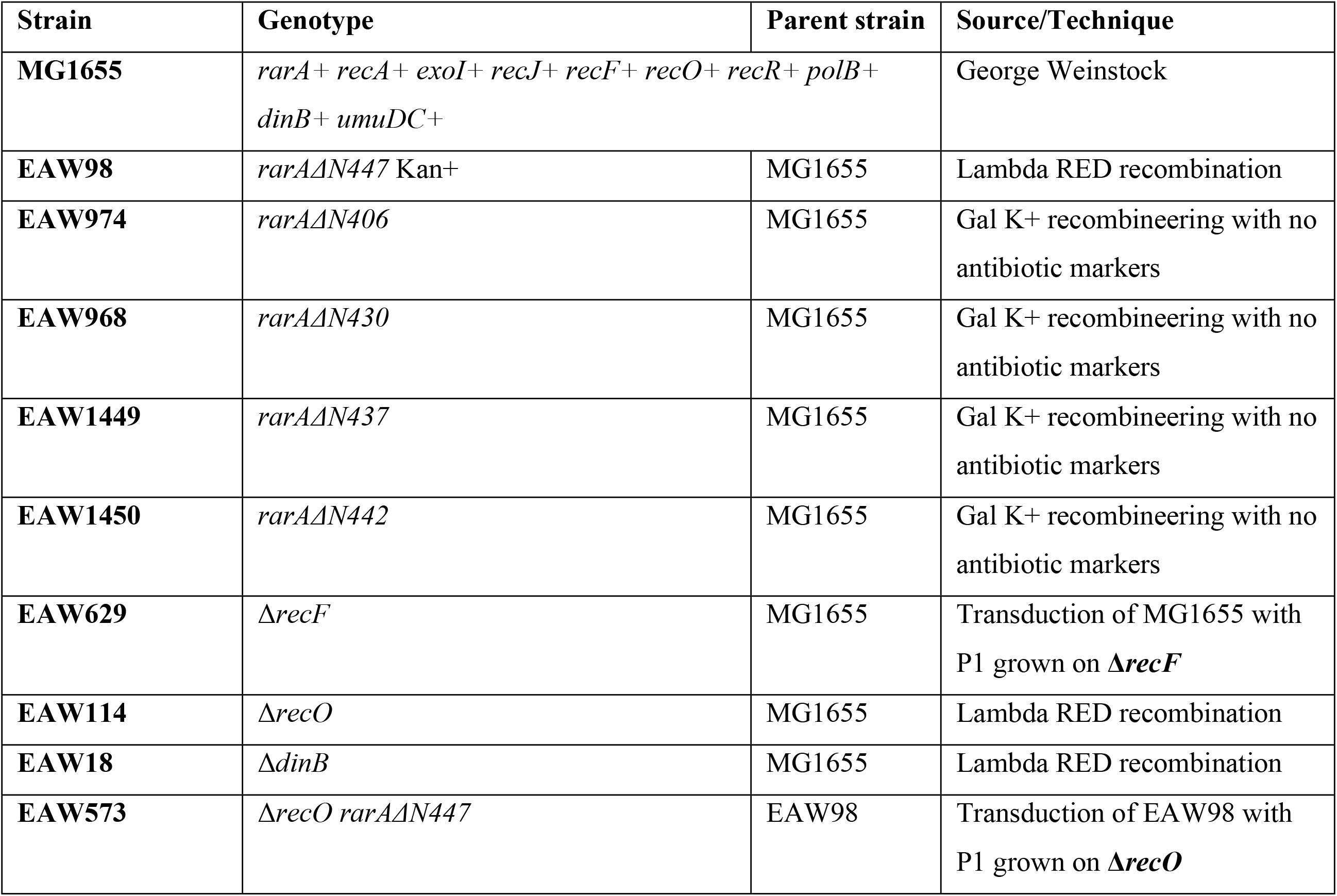
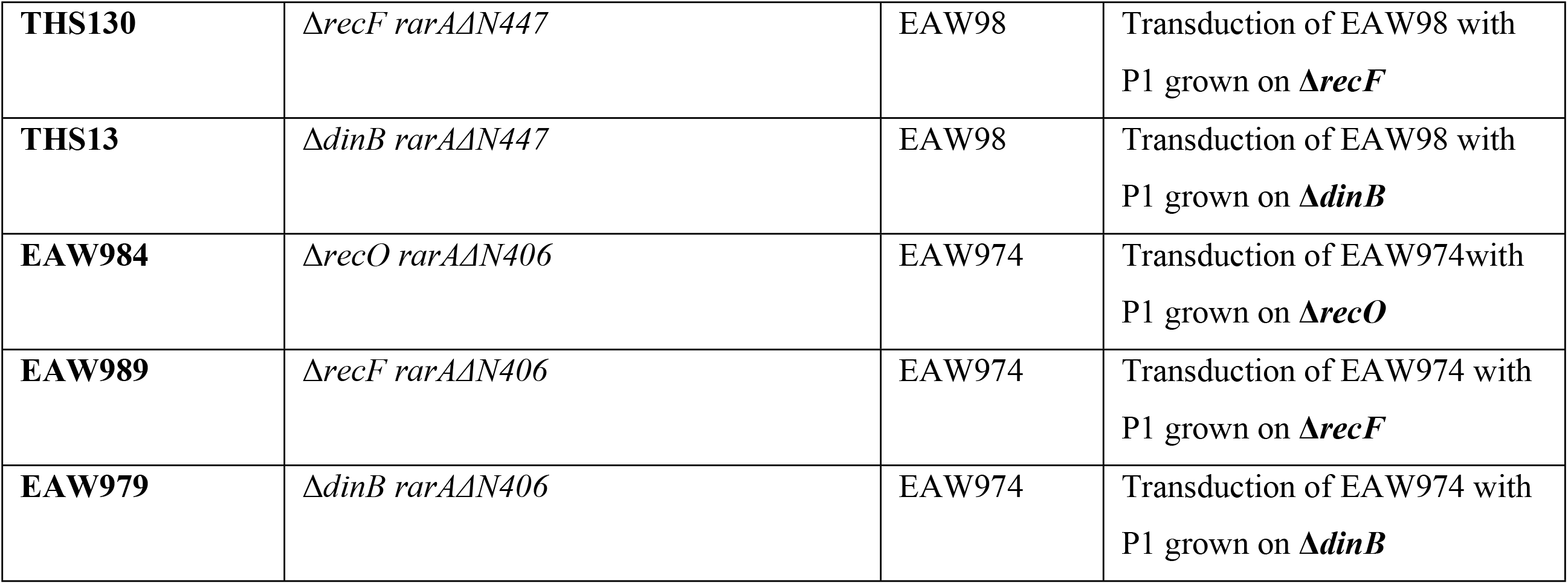
List of strains used in this study.

### Plasmid construction

All plasmids were sequenced to confirm the correct mutation(s)/insertion(s) following their construction. pBAD-*serS* is a pBAD/myc-His A Nde + wt *serS.* pEAW1176 was constructed by amplifying the wildtype *serS* gene containing NdeI and BamHI restriction cut sites from the *E. coli* MG1655 genome in a PCR. pBAD/myc-His A Nde was cut with NdeI and BglII enzymes (BglII creates compatible sticky ends with BamHI), while the PCR product was cut with NdeI and BamHI enzymes. The PCR product was ligated into the pBAD/myc-His A. pEAW1012 is a derivative of pRC7 plasmid (a lac+ mini-F low copy derivative of pFZY1) that expresses a WT copy of the *rarA* gene.

### Growth curve

Overnight cultures of indicated strains were diluted 1:100 in LB medium. 100 μl of each culture was poured in a clear bottom 96 well plate (Corning). Cultures were grown at 37° C with orbital shaking in a BioTek Synergy 2 plate reader. OD_600_ values were taken every 10 minutes for over the course of 800 minutes. OD_600_ values were normalized by subtracting out the OD_600_ value of only LB media. All growth curves represent averaged values from the three biological replicate experiments.

### Growth competition assays

Growth competition assays were conducted as previously described (63) using a method originally described by Lenski (60). The Δ*araBAD* ΔParaB marker was included on either wild type (MG1655) or mutant (Δ*rarA*) in separate experiments to control for any effect the marker may have had on cell fitness.

### Drug sensitivity assay

Overnight cultures of indicated strains were diluted 1:100 in fresh LB medium. Cultures were grown at 37°C with aeration and shaking until the OD_600_ measured 0.2. 1 mL aliquots were taken from each culture and were serially diluted in 1X PBS buffer (137 mM NaCl, 2.7 mM KCl, 10 mM Na_2_HPO_4_, 1.8 mM KH_2_PO_4_, 1 mM CaCl_2_ and 0.5 mM MgCl_2_) to 10^−6^. 10 μL of each dilution were spot plated on LB agar plates containing the indicated media. Plates were incubated overnight at 37°C and imaged the following day using a FOTO/Analyst Apprentice Digital Camera System (Fotodyne, Inc.). All experiments were conducted at least three times.

### SOS induction assay

Plasmid expressing the Green Fluorescent Protein (GFP) under the regulation of *recN* promoter (pEAW903) was used in this assay. First, either empty vector (pQBi63) or pEAW903 was transformed into the appropriate strains (WT, *rarA*ΔN447, or *rarAΔN406*) and the transformants were selected on Amp100 (100 μg/ml) plates. The transformants were then grown in 3ml of LB + Amp100 medium overnight at 37°C. The next day, the cultures were diluted in 1:1000 ratio in LB + Amp100 broth and 150μl of sample were poured into each well of a 96-well plate (Corning Incorporated/Costar) and put into the plate reader (Synergy H1 Hybrid Reader by BioTek). The samples were allowed to grow for 1000 mins at 37°C, with OD_600_ and GFP fluorescence (488/515nm) recorded at every 10 minutes. Relative fluorescence was calculated by normalizing the fluorescence reading to the OD_600_ of the culture.

### Analysis of cell shape: Bright field microscopy

All cells were grown overnight at 37°C and the saturated culture diluted in 1:100 ratio and grown in LB media till O.D. reaches 1.0. 200 μL of culture were then pelleted down, resuspended in 1XPBS buffer and incubated with 2 μl of FM-64 dye (0.33M) on ice for at least 30 mins. For imaging, 2μl of this mixture were loaded onto 0.16mm thick borosilicate glass made coverslips (Azer scientific) and sandwiched with 1% agarose gel pad. For all measurements of cell size and filamentation, wide-field microscopy was conducted on a STORM/TIRF inverted microscope ECLIPSE Ti-E (Nikon) (100× objective). Images using DIA and dsRed filters were collected on an ORCA Flash 4.0 Hamamatsu camera. A bright-field and dsRed image (at 100 ms and 50ms exposure respectively) were taken at multiple fields of view to determine the cell shape and length. For analysis, all images were imported into MicrobeJ, an ImageJ plugin, to outline cells. Selected cells were manually filtered for any outliers. All strains were imaged in triplicates and the cell size of each strain is averaged compiling each repeat.

## Results

It is well documented that RarA is involved in the maintenance of genome stability in cells, but its precise function and mechanism of action remains an enigma. To identify the phenotypic contribution of RarA in cells, we created a MG1655 derivative carrying a full deletion of the *rarA* allele. No growth or viability phenotype has previously been ascribed to strains with a *rarA* gene deletion (42, 43, 56–58). Previous work has focused on a modified *rarA* gene in which a chloramphenicol cassette has replaced either the first 600 nucleotides of the *rarA* gene (42, 43, 56, 57) or codons 113-349 (58), both in an *E. coli* AB1157 background. As most of our constructs are based on *E. coli* strain MG1655, we constructed a complete *rarA* gene deletion in the MG1655 background using Datsenko and Wanner method (59) (**Figure 2A**), and then studied the effect of this deletion on cell fitness.

**Figure 2:**
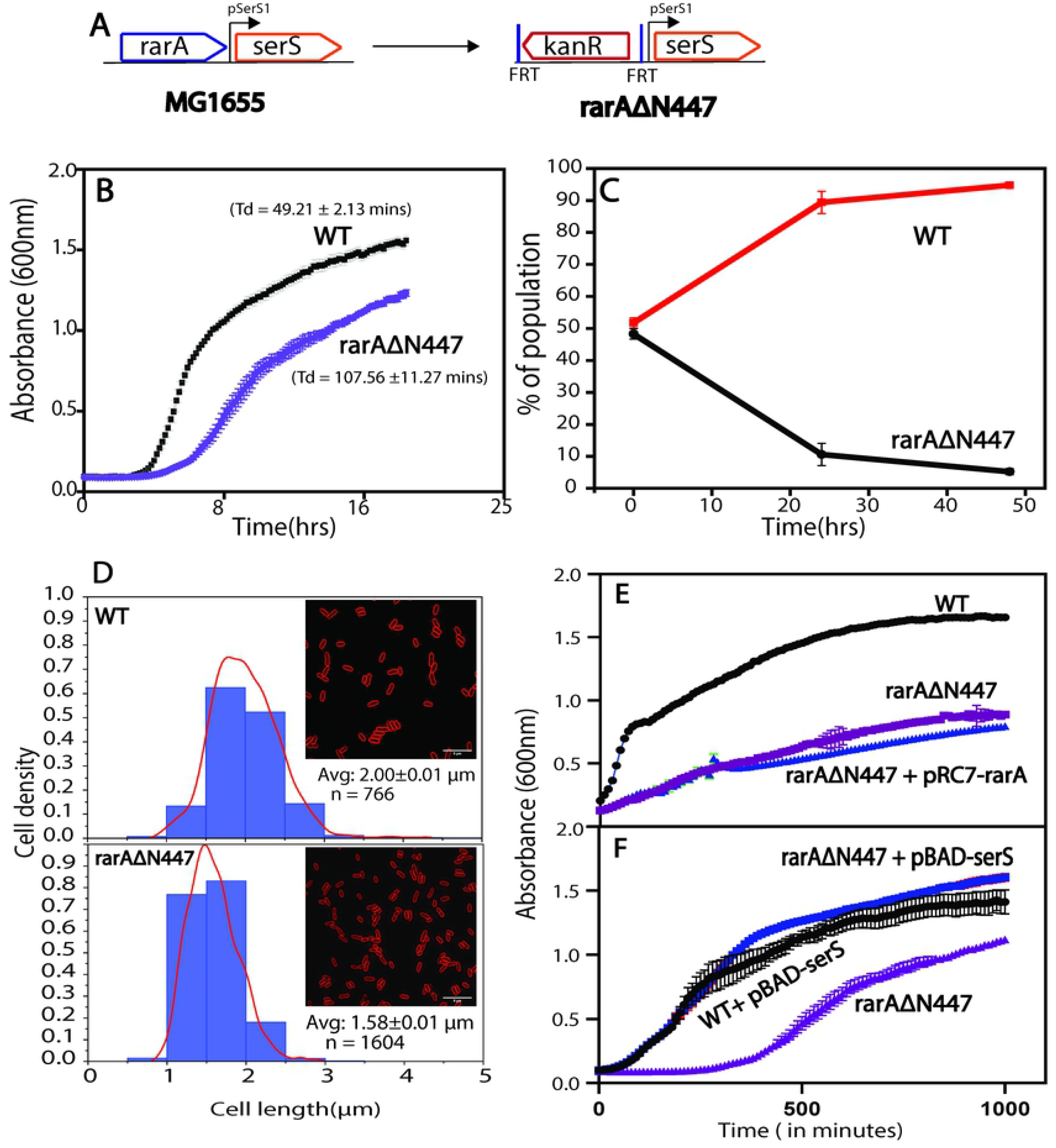
Complete deletion of *rarA (rarAΔN447)* causes a growth defect in *E. coli* cell. (A) Schematic of a complete deletion of *rarA* via a FLP recombinase method in *E. coli* chromosome. The *rarA* gene segment is replaced by a Kan cassette. (B) Growth curve: Deletion of complete *rarA* gene exhibit growth defect. (C) Growth competition: *rarAΔN447* is outcompeted by WT cells (D) Average cell size of *ΔrarA* cells decreases compared to WT cells. (E) Addition of pRC7-*rarA*, carrying WT *rarA* copy, in Δ*rarA* cell does not rescue its growth defect. (F) Overexpression of *serS* using pBAD vector rescues the growth defect of *ΔrarA* cells (blue line). Using the multi-genome browser of Ecocyc, we next searched for the orthologs of *rarA* in a broad range of organisms and then mapped the extent to which those orthologs have maintained their genetic context relative to *E. coli.* It revealed that the positioning of the *serS* gene - right downstream of the *rarA* gene locus - exists only in γ-proteobacteria class of the Proteobacteria phylum (**Figure 3**). Conservation of this proximity between *rarA* and *serS* genes across this class indicates their possible interconnection in other organisms of this class as well.

**Figure 3:**
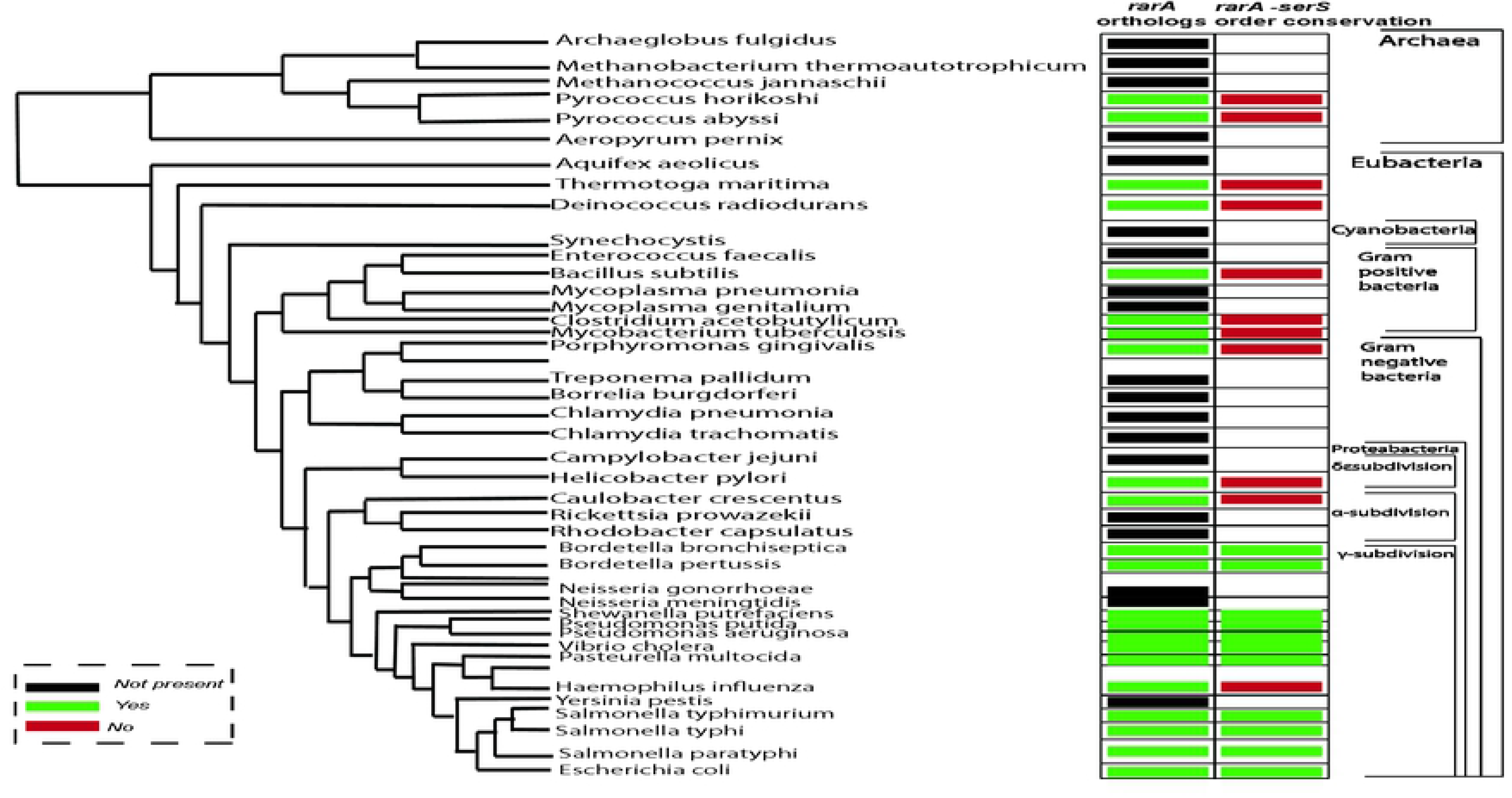
Multiple genome sequence alignment to identify orthologs of *rarA,* and the conservation of its genetic context in other organisms.

### Complete deletion of *rarA* causes a growth defect and reduced cell size of MG1655 *E. coli* cells

Using a plate reader, we noted and compared the growth of the *rarA*Δ*N447*strain to wild type cells at 37°C every 10 mins for 18hrs. The *rarA*Δ*N447* cells grew more slowly than wild type cells. (**Figure 2B**). To document the growth defect of the *rarA*Δ*N447* mutant with a different and more sensitive method, we carried out a direct competition assay between the wild type strain and the *rarA*Δ*N447*strain, using an approach developed by Lenski and colleagues (60). (**Figure 2C**). Wild type or mutant cells were modified to carry a neutral Ara– mutation (which confers a red color on colonies when grown on tetrazolium arabinose (TA) indicator plates) to permit color-based scoring of mixed populations. Overnight cultures of the *rarA*Δ*N447*strain were mixed in a 50/50 ratio with isogenic wild type cells carrying the Ara– mutation, or vice versa. The mixed culture was then diluted and grown up again on successive days, with plating to count red and white colonies occurring once each day. Earlier work (60, 61) demonstrated that the Ara– mutation does not affect growth rates by itself. We found that the wild type cells outgrew the *rarA*Δ*N447* cells and dominated the mixed cultures almost completely within 24 hours (**Figure 2C**). We further investigated the phenotypic dissimilarities between *rarA*Δ*N447* and WT cells using bright field microscopy. We observed that *rarA*Δ*N447* cells were substantially smaller than *rarA*+ cells (1.58μm [SEM =0.01] versus 2μm [SEM = 0.01] in length) (**Figure 2D**). RarA is well documented as a vital player in the DNA recombination and repair process. Suppressing the DNA repair system exerts stress response in the cell. With the data collected above, we presumed that the complete deletion of *rarA* impedes the damage tolerance capability of the cells that results in a significant growth impact.

### Phenotypic defects of *ΔrarA* cells are attributed to the lower expression of *serS* gene

For further affirmation of the results obtained above, we performed a complementation test. The pRC7 plasmid carrying a wild type copy of *rarA* along with ampicillin marker is employed in this study. We incorporated this plasmid into *rarA*Δ*N447* cell and test its growth rate. Surprisingly, we observed no rescue of the growth defect even in the presence of wild-type copy of *rarA* on the plasmid (**Figure 2E**). This observation signals that the growth defect observed earlier is not directly associated with the absence of *rarA* but might be an outcome of that deletion on other growth-related genes.

Based on the genomic location position of *rarA*, we hypothesized that the growth defect might be ascribed to the defect in the closest downstream essential gene - *serS*. It is well documented that the addition of SHX (serine hydroxamate), an inhibitor of the *serRS* gene, causes growth defects in *E. coli* cells even under the nutrient rich conditions (55). Changes in the levels of *serS* are expected to fluctuate the levels of charged to uncharged tRNA ratios and thereby the cell growth. Decreased cell viability of *rarA*Δ*N447* could be a result of the decreased *serS* levels in the cell. To test this hypothesis, we incorporated a plasmid overexpressing *serS* in *rarA*Δ*N447* cells. Overproduction of *serS* rescued the growth defect of *rarA*Δ*N447* cells (**Figure 2F**). This result signals the presence of promoter element/regulatory sequence for *serS* gene within the coding region of a *rarA* gene. Removal of that segment impacts the level of *serS* in the cell.

### Identification of promoter/regulatory sequence for *serS* gene in *rarA* sequence

We next aimed to identify the segment within a *rarA* gene that is controlling the *serS* expression under normal conditions. We constructed various *rarA* mutations differing in the number of nucleotides deleted from the N- terminus of the *rarA* to figure out the minimum region of *rarA* required to remain intact to mitigate the growth defect of *rarA*Δ*N447* cells. GalK+ recombineering method instead of Datensko Warner method was used to avoid any effect of kanamycin cassette sequence on the *serS* expression. We created four variants - *rarAΔ*N406, *rarAΔ*N430, *rarAΔ*N437, and *rarAΔ*N442, and studied their growth and cell morphology profiles (**Figure 4A).** Interestingly, none of these mutants showed any growth adversity like *rarAΔ*N447 (**Figure 4B and 4C**). However, the creation of complete deletion of *rarA* via galK recombineering method failed. This indicates that there exists a possible promoter or regulatory sequence for *serS* within the last 5 codons of *rarA –* deletion of which hampers the *serS* expression and thereby the growth of the cell. *serS* is mainly regulated by its promoter (serSp1) with a transcription start located at 28 nucleotides downstream from the end of *rarA* gene (46). Tracing back its possible promoter region, we suspected that the –10 region for this serSp1 is located at ~14 nucleotide from the *rarA* end and its –35 is located (although there is no good consensus –35 there) within the last ~15 nucleotides of *rarA* gene. A –35 segment of the *serS* promoter or some equivalent regulatory sequence within this last segment of *rarA* gene makes the complete deletion of *rarA* an infeasible option in the *E. coli* cell.

**Figure 4:**
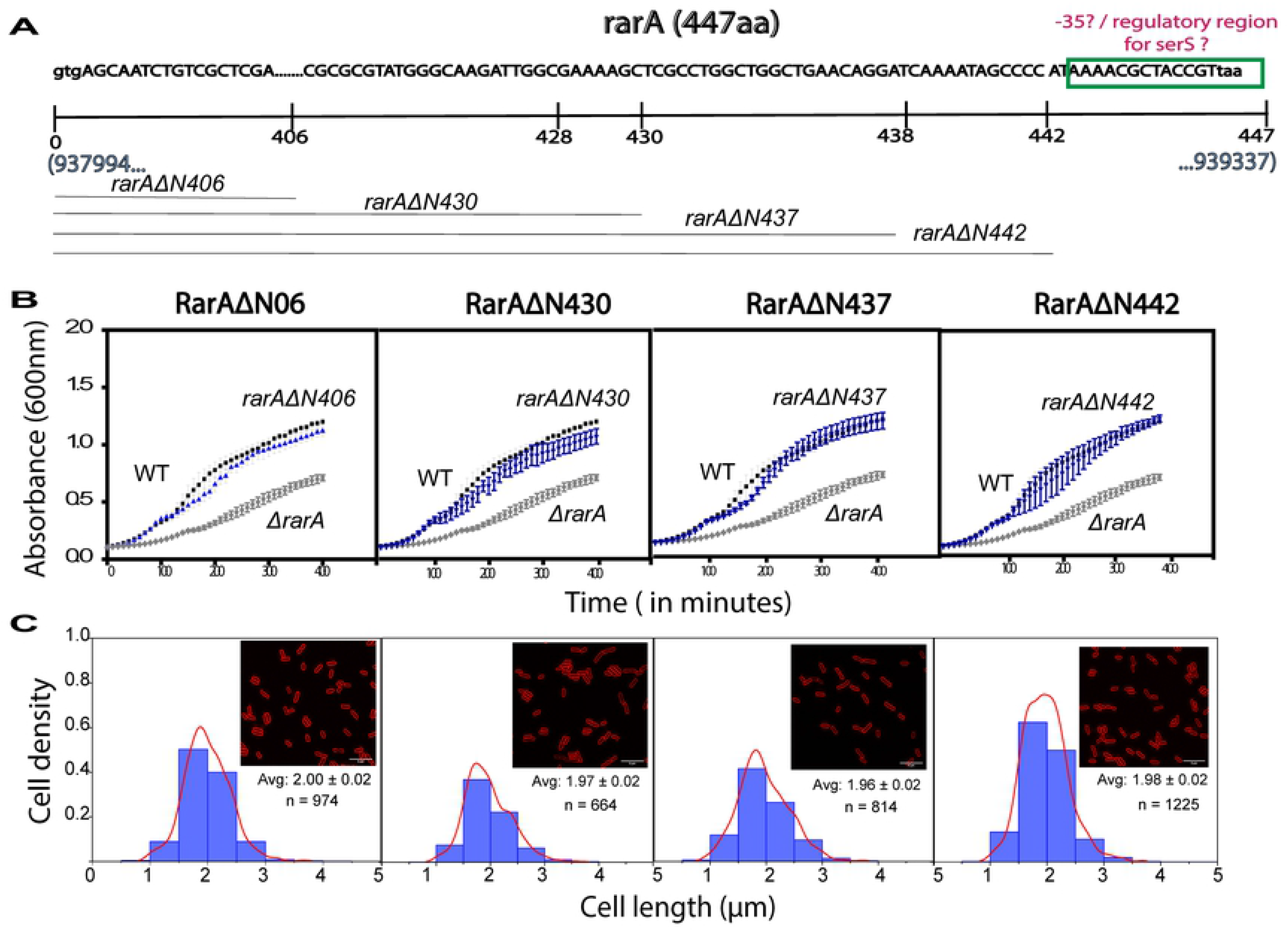
Analysis of the effect of various *rarA* deletion (*rarAΔN406, rarAΔN430, rarAΔN437, rarAΔN442, and rarAΔN447*) on the growth and cell size. (A) Schematic of *rarA* gene, highlighting the possible promoter regions and positions of different deletions made. (B) Growth curve: *ΔrarAN406*, *ΔrarAN430*, *ΔrarAN437, ΔrarAN442* does not exhibit a growth defect (C) Cell size: *rarAΔN406*, *rarAΔN430*, *rarAΔN437, rarAΔN442* has cell morphology comparable to WT.

### Consequences of Δ*rarA* on damage sensitivity of cells compromised with other DNA repair system

We tested the drug sensitivity of Δ*rarA447* cell alone and it in combination with other repair systems. The absence of a complete *rarA* sequence itself does not increase the cell sensitivity to DNA damaging agents like UV or NFZ. Removal of a RecA-loading system like RecF or RecO, however, increases the sensitivity to almost all kinds of damages. Interestingly, deletion of *rarA* in *recF* or *recO* background rescues their damage sensitivity. Complete deletion of*rarA* in *Δpol IV* background also decreases the sensitivity of *pol IV*-cells to both UV and NFZ induced damages (**Supplementary fig. 1)**. Moreover, we observed that SOS response is also induced in *rarA*ΔN447, at both with and without external damaging conditions (UV treatment). The induction was much higher than a WT cell (**Supplementary fig. 2)**. Interestingly, none of these results were replicated when Δ*rarA*N406 background was used instead of *rarA*ΔN447.

With all these observations, we confirmed that the phenotype due to the complete deletion of *rarA* is attributed to the decreased levels of *serS* in the cell. Decreased level of *serS* could cause a stringent response which activates the level of ppGpp. High levels of ppGpp act by rescuing the stalled RNA polymerases. The rescue of Δ*pol IV/ΔrecF/ΔrecO* cells’ drug sensitivity on deletion of *rarA*ΔN447 may be due to the rescue of stalled RNA polymerases, an outcome of the action of high levels of ppGpp in *rarA*ΔN447 cells. High SOS levels could also be a repercussion of this same phenomenon. Deletion of all but the last 41 codons of *rarA* eliminates all these phenotypes.

## Discussion

The major conclusion of this work is straightforward. Genetic elements affecting the expression of the *serS* gene are embedded in the final five codons of the upstream *rarA* gene. Upon complete deletion of *rarA*, we had documented a variety of phenotypic effects (supplementary data) that we initially attributed to a loss of *rarA* function. These disappeared when we made use of *rarA* deletions that encompasses most but not all of the gene. We now attribute the effects to changes in *serS* expression, possibly reflecting some aspect of a stringent response.

The *serS* promoter element that is within the *rarA* gene has not been identified. The region in question is positioned so as to potentially include a –35 region for the promoter. However, no – 35 consensus is evident.

## Supporting information

**Supplementary Fig 1: Consequence of complete deletion of *rarA* on DNA damage sensitivity of the cell.** (A) Complete deletion of *rarA* is able to rescue the sensitivity of *ΔrecF* and *ΔrecO* to UV and *ΔdinB* to NFZ. (B) Incorporation of *ΔrarAN406* does not rescue the sensitivity of *ΔrecF* and *ΔrecO* to UV and *ΔdinB* to NFZ.

**Supplementary Fig 2: SOS induction is highly induced on UV exposure in cells carrying complete deletion of *rarA* (*rarAΔN447)* from the cell.** Complete deletion of *rarA* induces SOS response more than WT cell, in presence and absence of UV exposure.

